# Synthetic Frizzled agonist and LRP antagonist for high-efficiency Wnt/β-catenin signaling manipulation in organoid cultures and *in vivo*

**DOI:** 10.1101/2023.06.21.545860

**Authors:** Quanhui Dai, Jiawen Wang, Zihuan Lin, Danni Yu, Hui Yang, Jiting Zhang, Xiaoyu Li, Junwei Zeng, Hao Hu, Chao Ni, Bing Zhao

## Abstract

Wnt/β-catenin signaling and its dysregulation play critical roles in the fate determination of stem cells and the pathology of various diseases. However, the application of translated Wnt ligand in regenerative medicine is hampered by its hydrophobicity and cross-reactivity with Frizzled (FZD) receptors. Here, we generate an engineered water-soluble, FZD subtype-specific agonist, RRP-pbFn, for high-efficiency Wnt/β-catenin signaling activation. In the absence of direct binding to LRP5/6, RRP-pbFn stimulates Wnt/β-catenin signaling more potently than surrogate Wnt. RRP-pbFn supports the growth of a variety of mouse and human organoids, and induces the expansion of liver and intestine progenitors *in vivo*. Meanwhile, we develop a synthetic LRP antagonist, RRP-Dkk1c, which exhibits heightened effectiveness in attenuating Wnt/β-catenin signaling activity compared to Dkk1, thereby abolishing the formation of CT26-derived colon cancer xenograft *in vivo*. Together, these two paired Wnt/β-catenin signaling manipulators hold great promise for biomedical research and potential therapeutics.

## Introduction

The Wnt/β-catenin signaling tightly controls embryonic development and tissue homeostasis by regulating stem cell self-renewal and lineage specification.^1, 2^ Wnt ligands heterodimerize receptors Frizzled (FZD) and low-density lipoprotein receptor-related proteins 5/6 (LRP5/6), which triggers the cytoplasm accumulation of core effector β-catenin then its nuclear translocation for target genes transcription.^3, 4^ The dysregulation of Wnt/β-catenin signaling is involved in the pathological processes of various diseases, encompassing developmental defects, cancers, and degenerative disorders.^1, 5, 6^ Therefore, manipulation of Wnt/β-catenin signaling activity *in vivo* would permit the development of novel therapeutic strategies.

*In vitro* stem cell expansion and programed differentiation bring the dawn of regenerative medicine. Since Wnt/β-catenin signaling governs stem cell self-renewal, its agonists have been utilized in maintaining stem/progenitor cell expansion. Especially, the newly emerging organoids, derived from adult stem cells through controlled fate determination, have shown great advantages in modeling various development events and diseases for drug discovery, precision medicine and regenerative theraputics.^7–9^ The design of more potent and stable Wnt manipulators will significantly advance stem cell-related technologies, particularly organoid cultures.

Extracellular signal ligands bind to specific cell surface receptors to initiate signaling transduction. Therefore, signal receptor agonist exerting the function of natural ligands or antagonist enables the precise regulation of signaling cascades without penetrating cell membrane, which presents great advantages in specificity, safety and delivery compared to intracellular targeting strategies.^10^ For Wnt/β-catenin signaling, intracellular agonists bear a greater risk of off-target effects, such as glycogen synthase kinase-3 (GSK3) inhibitor CHIR exerting regulatory effects on diverse cellular pathways beyond Wnt/β-catenin signaling and leading to much broader alterations in global gene expression compared to Wnt3a ligand or its surrogate.^11^

However, the promiscuous interactions arising from the conserved interaction sites between the 19 mammalian Wnt ligands and the 10 FZD subtypes (FZD 1-10) have hindered in unequivocally assigning distinct downstream responses to individual FZD subtypes.^12^ Furthermore, Wnt ligands undergo post-translational palmitoylation facilitated by the O-acyltransferase known as Porcupine, which is essential for both Wnt secretion and interaction with FZD.^13, 14^ As a result of their palmitoleate groups, Wnt ligands exhibit high hydrophobicity, which necessitates the utilization of detergents for purification and consequently poses challenges in purifying and employing recombinant Wnt ligands.^14^ Wnt3a conditioned media (CM) is commonly employed as Wnt/β-catenin signaling activators. Nevertheless, the exact composition of Wnt3a CM is not fully determined, introducing uncontrollable variables in experimental settings.^15^ Additionally, FZD subtype-specific surrogate Wnt (sWnt), which involve linking the protein binder of FZD (pbF, also known as DRPB, binding to FZD1, 2, 5, 7, 8) to the C-terminal domain of the human Wnt antagonist Dkk1 (Dkk1c), as well as tetrameric antibody-based Wnt agonists, have been reported as macromolecules.^16–19^ These molecules induce heterodimerization or oligomerization of FZD-LRP, activating the Wnt/β-catenin signaling and bypassing the need for endogenous Wnt ligands. The challenges associated with manufacturing the endogenous ligands and large molecular size of currently available Wnt agonists render them impractical as viable drug targets. Therefore, it is imperative to engineer water-soluble, high-efficiency, and FZD-specific synthetic agonists to elucidate the intricate mechanism of Wnt/β-catenin signaling activation and facilitate organoid generation.

Inappropriate increase in Wnt receptor activity due to mutational inactivation of the ubiquitin ligases RNF43/ZNRF3 or R-spondin fusions has recently been identified as a prominent driver of cancer development.^20–22^ The antagonists targeting canonical Wnt receptors hold promise for treating these specific cancer subsets.^23–25^ Thus, the development of synthetic LRP antagonists has great prospects for therapeutic applications.

Here, we report the development of two synthetic water-soluble Wnt manipulators: the FZD agonist RRP-pbFn, which consists of RRP (truncation of optogenetic tool pMag) fused with pbFn (N-terminal domain of pbF),^26–28^ and the LRP antagonist RRP-Dkk1c. Compared to the improved surrogate Wnt (Dkk1c-pbF), RRP-pbFn exhibits distinct characteristics, as it does not directly bind to the LRP5/6 receptor, possesses a lower molecular weight, and activates Wnt/β-catenin signaling more potently. RRP-pbFn demonstrates potent support for diverse types of organoid growth and expansion, including small intestine, alveolus, stomach, hepatocyte, and cholangiocyte. Furthermore, we demonstrate the *in vivo* efficacy of RRP-pbFn in regulating metabolic liver zonation and promoting adult intestinal proliferation. Synthetic RRP-Dkk1c manifests superior efficacy in Wnt/β-catenin signaling inhibition compared to Dkk1, resulting in the inhibition of organoid growth and effective suppression of tumor growth in the CT26 xenograft model. These easily produced water-soluble Wnt/β-catenin signaling manipulators hold significant potential for exploring research and translational applications in regenerative medicine.

## Results

### Generation of a synthetic Frizzled agonist for high-efficiency Wnt/β-catenin signaling activation

The precise spatial and temporal control of cell signaling has widespread application in regenerative medicine and the study of tissue development.^29, 30^ We initially aimed to develop a photoswitch-controlled sWnt for spatiotemporal control of Wnt/β-catenin signaling. We employed the light-activated Magnets dimerization system by fusing different orientations of the sWnt fragments (pbF/Dkk1c) with the pMag/nMag (Figure S1A).^16, 26^ Variants of sWnt/Magnets halves were transfected individually or in combination into human embryonic kidney 293T (HEK 293T) cells along with TOP-Flash and *Renilla* reporters. The luciferase activity was subsequently determined with or without illumination. However, the pMag/nMag-based sWnt system showed no enhanced Wnt activity after illumination, suggesting that splitting sWnt into pbF and Dkk1c was unsuitable for Magnets photoswitch-controlled sWnt activity (Figure S1B). Surprisingly, pMag-pbF alone, even without a combination or blue light, induced significant and robust Wnt activity (Figure S1B).

Following the observation of potent Wnt activity elicited by the pMag-pbF fusion protein, we subsequently tested the capacity of pMag and pbF individually or collaboratively, to activate the Wnt/β-catenin signaling. As a consequence, only the pMag-pbF fusion protein showed a 29.8-fold inducible effect on Wnt activity, whereas neither pMag alone, pbF alone, nor their combination exhibited such effect (Figure 1A). 50% pMag-pbF conditioned media (CM) also activated the Wnt/β-catenin signaling (Figure 1B). Interestingly, we observed that orientation had an impact on activity for pMag-pbF fusion protein, where pbF-pMag (pbF fused to the N-terminal of pMag) exhibited lower activity than pMag-pbF (pMag fused to the N-terminal of pbF) in inducing Wnt activity (Figure 1C). In addition, pMag-pbF showed synergy with Wnt potentiator R-spondin1,^31^ demonstrated by the enhanced expression of the TOP-Flash reporter in HEK 293T cells (Figure 1D). The substitution of the amino acid residue Ile52 and Met55 of VVD with R/R (I52R/M55R) or D/G (I52D/M55G) gave rise to pMag and nMag, respectively.^26^ Subsequently, we investigated the capacity of the nMag-pbF and VVD-pbF fusion proteins to induce Wnt/β-catenin signaling, but no activation was observed (Figure 1E). Structural analysis of pMag, VVD, and nMag proteins revealed that structural variations were restricted to specific amino acid residues 52 and 55 (Figure 1F). Based on this observation, we hypothesized that R52 and R55 residues of pMag were critical for the agonist activity of pMag-pbF. To confirm this, we generated pMag variants with substitutions of R52 with I or R55 with M, resulting in pMag(R52I) and pMag(R55M), respectively (Figure 1F), which were subsequently fused with pbF. Our findings confirmed the hypothesis, as pMag(R55M)-pbF showed decreased activity, whereas pMag(R52I)-pbF had no activity when compared to pMag-pbF (Figure 1E).

**Figure 1.**
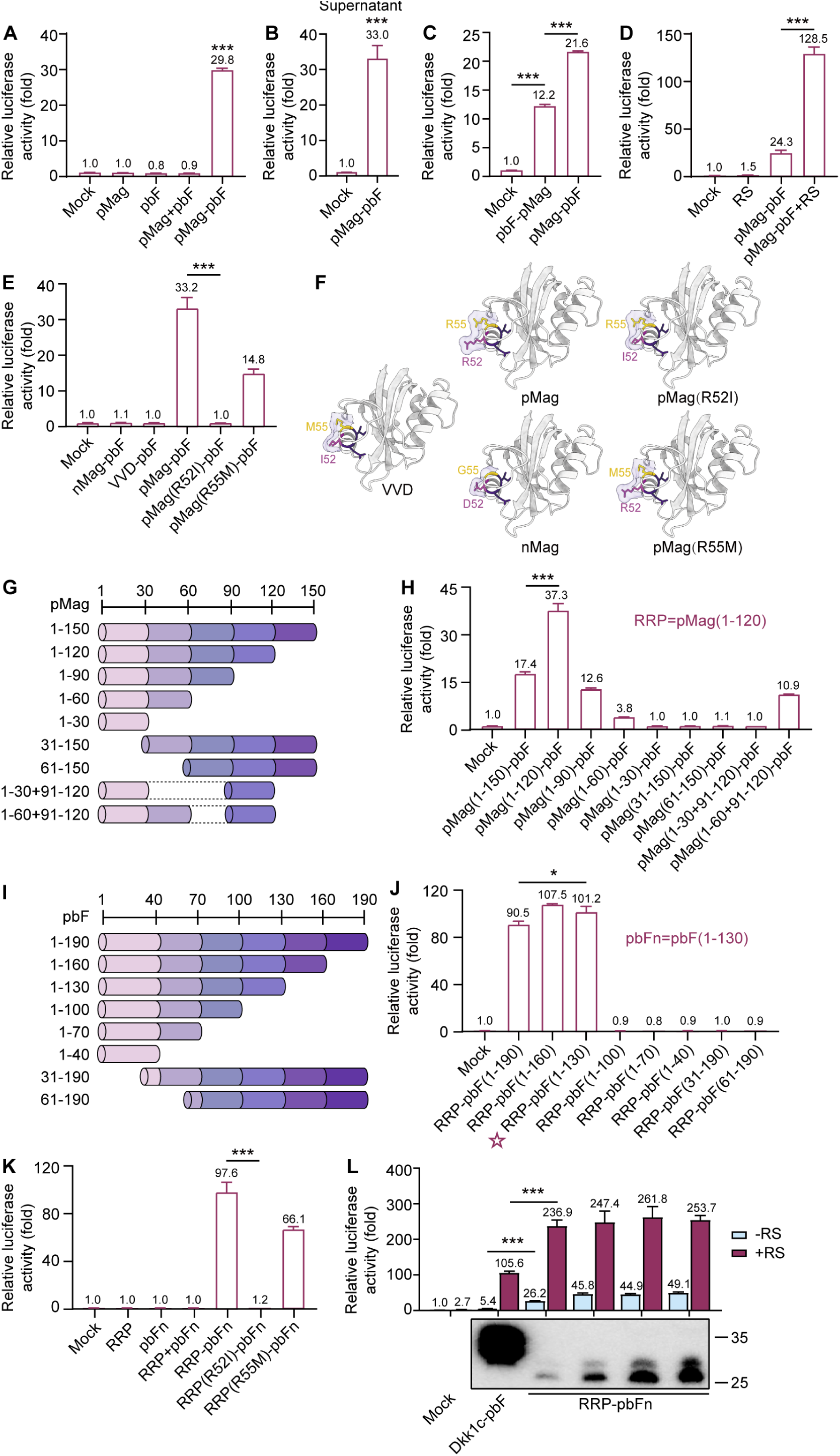
Generation of a synthetic Frizzled agonist for high-efficiency Wnt/β-catenin signaling activation. (A) TOP-Flash luciferase reporter assay showing the Wnt activity induced by overexpression of pMag-pbF or control proteins. The empty vector (mock) along with reporter plasmids was used as a negative control. (B) TOP-Flash luciferase reporter assay showing activation of Wnt/β-catenin signaling by 50% pMag-pbF CM. (C) TOP-Flash luciferase reporter assay showing the Wnt activity induced by overexpression of pMag-pbF (pMag fused to the N-terminal of pbF) and pbF-pMag (pbF fused to the N-terminal of pMag). (D) 250 ng/mL R-spondin1 potentiated the Wnt activity induced by pMag-pbF overexpression. (E) The Wnt activity induced by nMag-pbF, VVD-pbF, pMag(R52I)-pbF, and pMag(R55M)-pbF overexpression. (F) The structure of VVD, pMag, nMag, pMag(R52I), and pMag(R55M) with different amino acid residues. VVD has amino acids I/M at positions 52 and 55, while pMag and nMag have R/R and D/G, respectively. Mutated pMag(R52I) and pMag(R55M) have I/R and R/M, respectively. Surface of each protein is showed as light purple patches. (G) Schematic representation of the full-length and various truncated pMag. The numbers listed on the left represent the position of the site of truncation based on a full-length pMag. (H) The Wnt activity induced by overexpression of the variants of full-length or truncated pMag fused with pbF. RRP = pMag(1-120). (I) Schematic representation of the full-length and various truncated pbF. The numbers listed on the left represent the position of the site of truncation based on a full-length pbF. (J) The Wnt activity induced by overexpression of RRP fused with the variants of full-length or truncated pbF. pbFn = pbF(1-130). (K) The Wnt activity induced by overexpression of RRP, pbFn, RRP+pbFn, RRP-pbFn, and mutated RRP(R52I)-pbFn and RRP(R55M)-pbFn. (L) Overexpressing RRP-pbFn induced higher dose-dependent Wnt/β-catenin signaling activation than the improved surrogate Wnt (Dkk1c-pbF) with or without 250 ng/mL R-spondin1. The bottom panel is detection of the improved surrogate Wnt (Dkk1c-pbF) and RRP-pbFn in supernatant by Western blot after overexpression. All TOP-Flash data represent mean ± SD; *n* = 3 independent experiments. \**p* < 0.05; \*\**p* < 0.01; \*\*\**p* < 0.001.

The utility and druggability of synthetic agonists depend heavily on the molecular weight. We then explored the shortest variant with highest-efficiency Wnt/β-catenin signaling activation by truncating pMag and pbF separately. Truncated variants of pMag (150 aa in total) and pbF (190 aa in total) were generated by stepwise deleting 30 aa from either the N or C-terminal (Figures 1G and 1I). Following fusion with pbF, pMag(1-120)-pbF exhibited the strongest Wnt activity, while pMag(1-90)-pbF showed decreased activity compared to pMag-pbF (Figure 1H). Although pMag(1-60)-pbF showed weak activity, pMag(1-60+91-120)-pbF restored the activity. Thus, we named pMag(1-120) as RRP. Subsequently, following fusion with RRP, RRP-pbF(1-130) displayed more potent activation than RRP-pbF(1-190) (Figure 1J), and pbF(1-130) was named pbFn. Furthermore, RRP-pbFn-induced Wnt activity remained unchanged after blue light illumination (Figure S2A). Using mutagenesis of RRP within pMag(R52I) and pMag(R55M), we observed that RRP(R55M)-pbFn displayed diminished activity, whereas RRP(R52I)-pbFn showed no activity, consistent with the behavior of pMag-pbF (Figure 1K). Moreover, pbFn and RRP(R52I)-pbFn inhibited the activation of Wnt/β-catenin signaling induced by RRP-pbFn (Figure S2B).

### RRP-pbFn activates Wnt/β-catenin signaling more potently than improved surrogate Wnt

We then set out to determine whether FZD agonist RRP-pbFn exhibits superior capacity for activating Wnt/β-catenin signaling. Previously reported sWnt (pbF-Dkk1c) induced Wnt/β-catenin signaling activity,^16^ and when the LRP6 binder was fused to the N-terminus of the FZD binder resulting in higher activity in comparison with the other orientation.^17^ Therefore, we compared the activity of improved surrogate Wnt (Dkk1c-pbF) with RRP-pbFn in a dose-dependent manner using Wnt reporter assays in HEK 293T cells. RRP-pbFn induced dose-response and stronger Wnt/β-catenin signaling even at low concentration than improved surrogate Wnt with or without R-spondin1 (Figure 1L). Additionally, RRP-pbFn exhibited a smaller molecular weight in comparison to improved surrogate Wnt (Figure 1L). In summary, RRP-pbFn, a water-soluble synthetic FZD agonist with a smaller molecular weight, activates Wnt/β-catenin signaling even much more potently than improved surrogate Wnt. These results suggest that targeting FZD might be a promising strategy to discover next-generation Wnt agonist.

### RRP-pbFn facilitates the growth of various mouse and human organoids

As Wnt/β-catenin signaling plays a dominant role in tissue stem cell self-renewal thus long-term maintenance *in vitro*, its ligand or surrogate is widely employed in the culture system of stem/progenitor cells even organoids.^16, 32, 33^ We evaluated whether RRP-pbFn would act as a next-generation Wnt/β-catenin sustainer for organoid cultures. Multiple types of organoids were generated from mouse small intestine, alveolus, stomach, hepatocyte, and human cholangiocyte, then cultured in the presence or absence of recombinant RRP-pbFn. The basal media-cultured intestinal organoids displayed normal morphology as previously described,^33^ whereas RRP-pbFn-supported organoids became thin-walled cystic structures, indicating more proliferation (Figure 2A). RRP-pbFn demonstrated tremendous improvement of the number of stomach-derived organoids and the area of alveolus-, hepatocyte-, and human cholangiocyte-derived organoids (Figures 2B-2E). Moreover, this was strongly paralleled by the enhanced transcription level of Wnt target genes *Lgr5* and *Axin2* by RRP-pbFn (Figure 2F). Collectively, these findings demonstrate that RRP-pbFn supports the growth of various organoid models.

**Figure 2.**
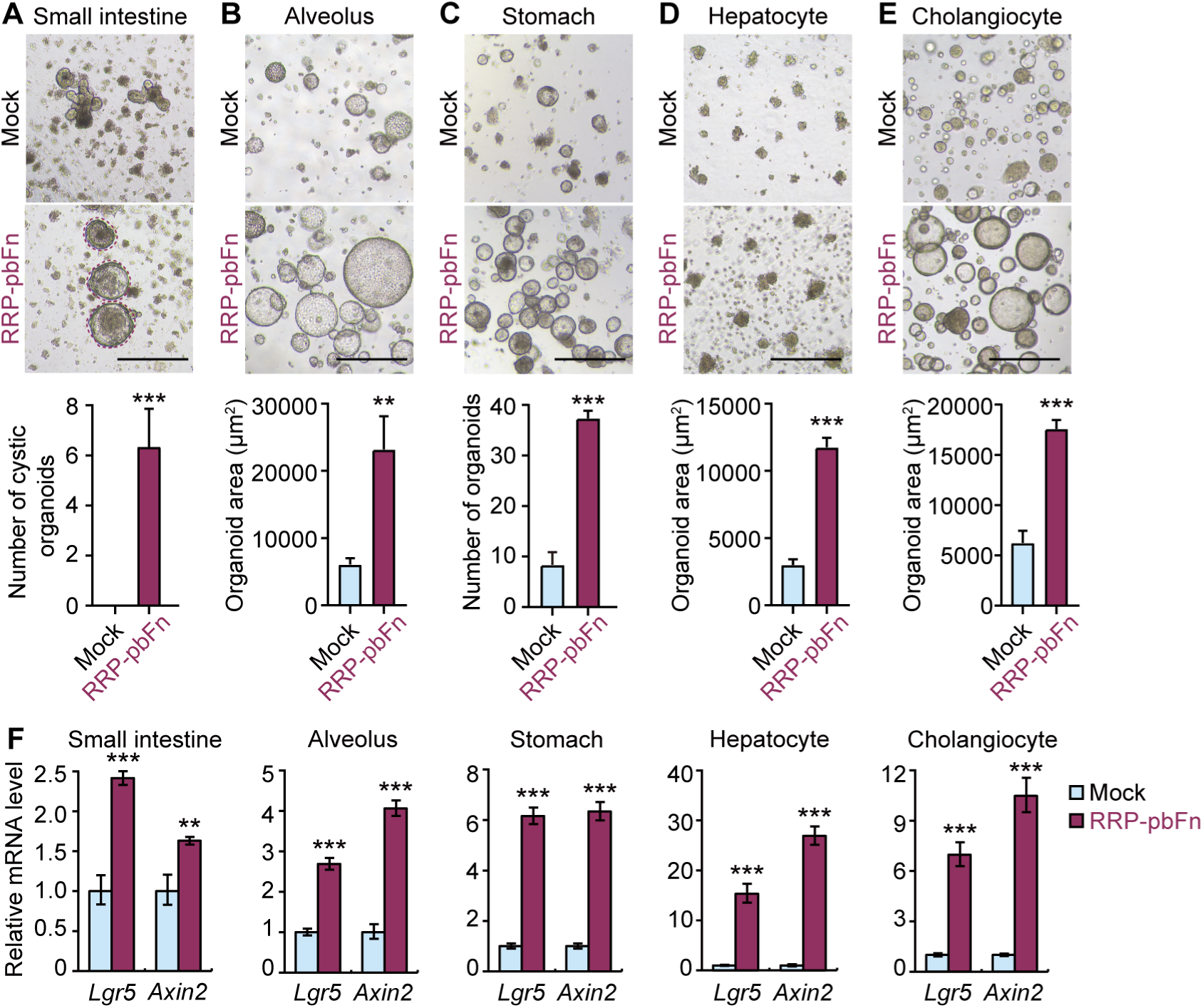
RRP-pbFn facilitates the growth of various mouse and human organoids. (A-E) Representative bright-field images of mouse organoids derived from small intestine (A), alveolus (B), stomach (C), and hepatocyte (D) expanded in culture media, and human cholangiocyte (E) expanded in culture media containing 3 μM IWP-2, and supplemented with or without recombinant RRP-pbFn. The bottom panels are quantification of the organoid number (small intestine, stomach) or organoid area (alveolus, hepatocyte, and cholangiocyte). Data represent mean ± SD, and *n* = 3 for each organoid type. The scale bar represents 500 μm. (F) qRT-PCR analysis of gene expression of Wnt/β-catenin signaling target genes. Data represent mean ± SD. \**p* < 0.05; \*\**p* < 0.01; \*\*\**p* < 0.001.

### RRP-pbFn induces the expansion of liver and intestine progenitors through manipulating Wnt/β-catenin signaling *in vivo*

As FZD agonist RRP-pbFn exhibits high-efficiency and stable ability of regulating cell fate determination in the near-physiology organoids, we then moved to testify whether RRP-pbFn works *in vivo*. The physiological processes of adult liver zonation and intestinal stem cell expansion represent archetypal Wnt-responsive events.^1, 16, 34^ To assess the *in vivo* activity of RRP-pbFn, we utilized adeno-associated virus 8 (AAV-8), a delivery system that can infect hepatocytes in liver.^35^ RRP-pbFn was fused with mouse IgG Fc fragment (Fc) to prolong the circulating half-life. Subsequently, mice were injected with AAV-8 expressing RRP-pbFn-Fc or Fc as a negative control, followed by injection with or without recombinant R-spondin1 (Figure S3A). Validation of AAV-mediated RRP-pbFn-Fc expression in mouse liver was confirmed by qRT-PCR analysis (Figure S3B). In the liver, Wnt ligands are expressed in the vicinity of the central vein (CV) and the activation of Wnt/β-catenin signaling triggers the pericentral gene expression program in adjacent hepatocytes,^36^ including *glutamine synthetase* (*GS*), consequently perturbing metabolic zonation (Figure 3A). We observed that single treatment with AAV-RRP-pbFn-Fc did not significantly affect liver GS expression, but combinatorial treatment with recombinant R-spondin1 induced a synergistic expansion of the GS-expressing pericentral region (Figures 3B and 3C), and up-regulated *GS* and *Axin2* while repressing the periportal marker *Cyp2f2* mRNA compared with control mice (Figure 3D).

**Figure 3.**
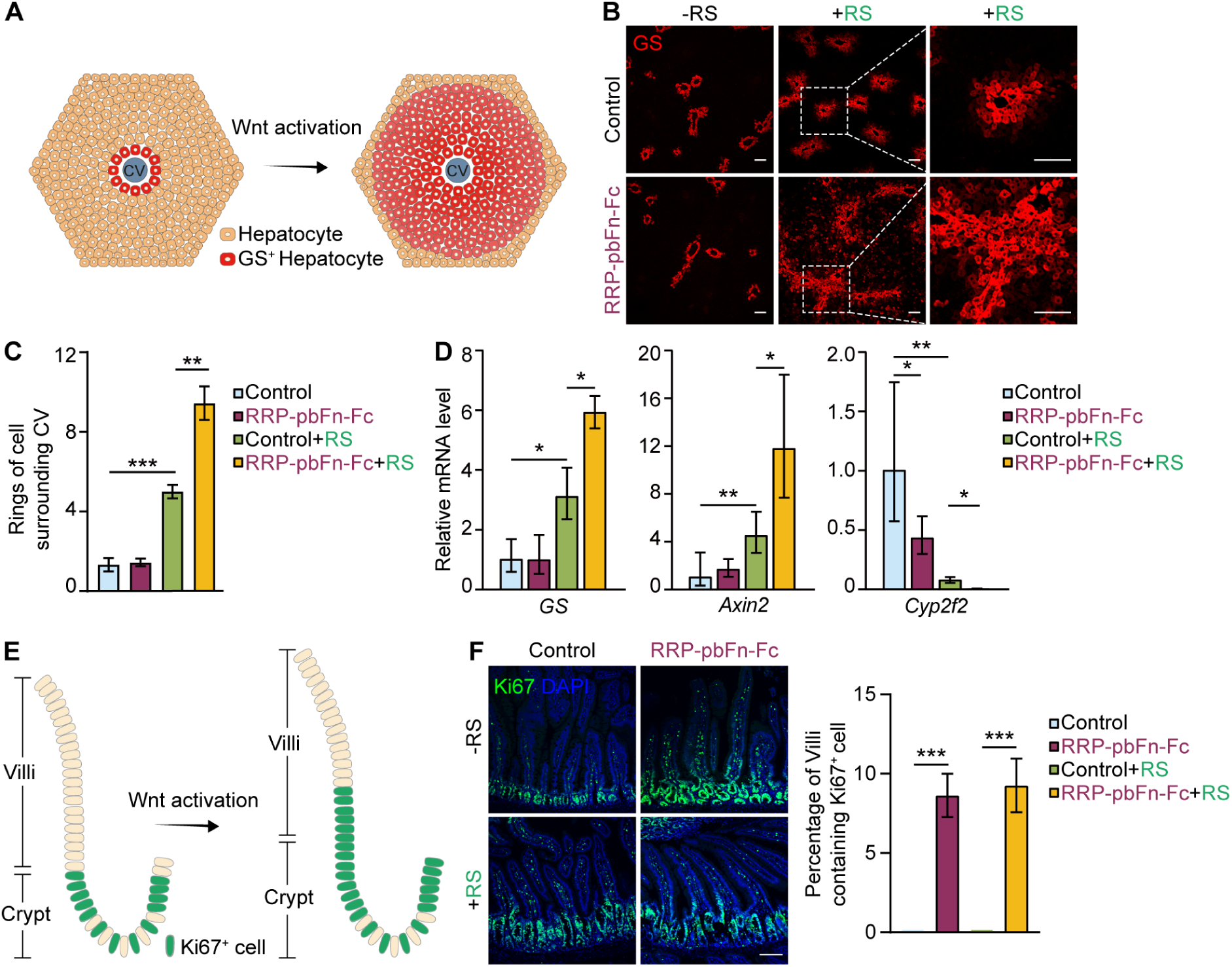
RRP-pbFn induces the expansion of liver and intestine progenitors through manipulating Wnt/β-catenin signaling *in vivo*. (A) Scheme depicting activation of Wnt/β-catenin signaling regulating liver zonation. (B) Representative images of GS (pericentral marker) immunofluorescence staining of livers from mice following AAV and R-spondin1 administration. RS: R-spondin1. The scale bar represents 100 μm. (C) Quantification of the rings of GS^+^ hepatocytes around the central vein. (D) qRT-PCR analysis of Wnt target genes *GS* and *Axin2*, periportal marker *Cyp2f2* of livers from mice received AAV and R-spondin1 administration. (E) Scheme depicting activation of Wnt/β-catenin signaling promoting adult intestinal proliferation. (F) Representative images of Ki67 immunofluorescence staining of jejunum from mice following AAV and R-spondin1 administration. DAPI staining is represented in blue. The scale bar represents 100 μm. The right panel is quantification of percentage of Villi containing Ki67^+^ cells. Data represent mean ± SD. *n* = 3 mice per group. \**p* < 0.05; \*\**p* < 0.01; \*\*\**p* < 0.001.

In the small intestine, Wnt/β-catenin signaling activation stimulates crypt hyperplasia and expansion of mitotic Ki67^+^ epithelium (Figure 3E). Upon intravenous injection, AAV-8 was capable of infecting mouse liver hepatocytes, resulting in persistent secretion of transgene products into the circulation. AAV-RRP-pbFn-Fc infection modestly induced Ki67 expression from the crypt extending toward the bottom region of Villi (Figure 3F). Hence, our results confirm that RRP-pbFn activates Wnt/β-catenin signaling *in vivo*, altering metabolic liver zonation and inducing intestinal villi proliferation.

### RRP-pbFn does not bind to LRP, but triggers FZD-LRP interaction for Wnt/β-catenin signaling activation

The activation of Wnt/β-catenin signaling occurs when the Wnt ligand binds to the coreceptors FZD and LRP5/6, leading to the dimerization of two receptors.^1^ It was known that Wnt/β-catenin signaling activation by RRP-pbFn was dependent on the recruitment of FZD receptor but unclear about LRP5/6 receptor. In order to determine whether Wnt/β-catenin signaling activation by RRP-pbFn binds directly to the LRP5/6 receptor, we conducted co-immunoprecipitation (Co-IP) in HEK 293T cells. The outcomes revealed that LRP5E1E4 and LRP6E1E4 Co-IP with sWnt, but not with RRP-pbFn (Figure 4A), suggesting that the activation of Wnt/β-catenin signaling by RRP-pbFn without binding to the LRP5/6 receptor.

**Figure 4.**
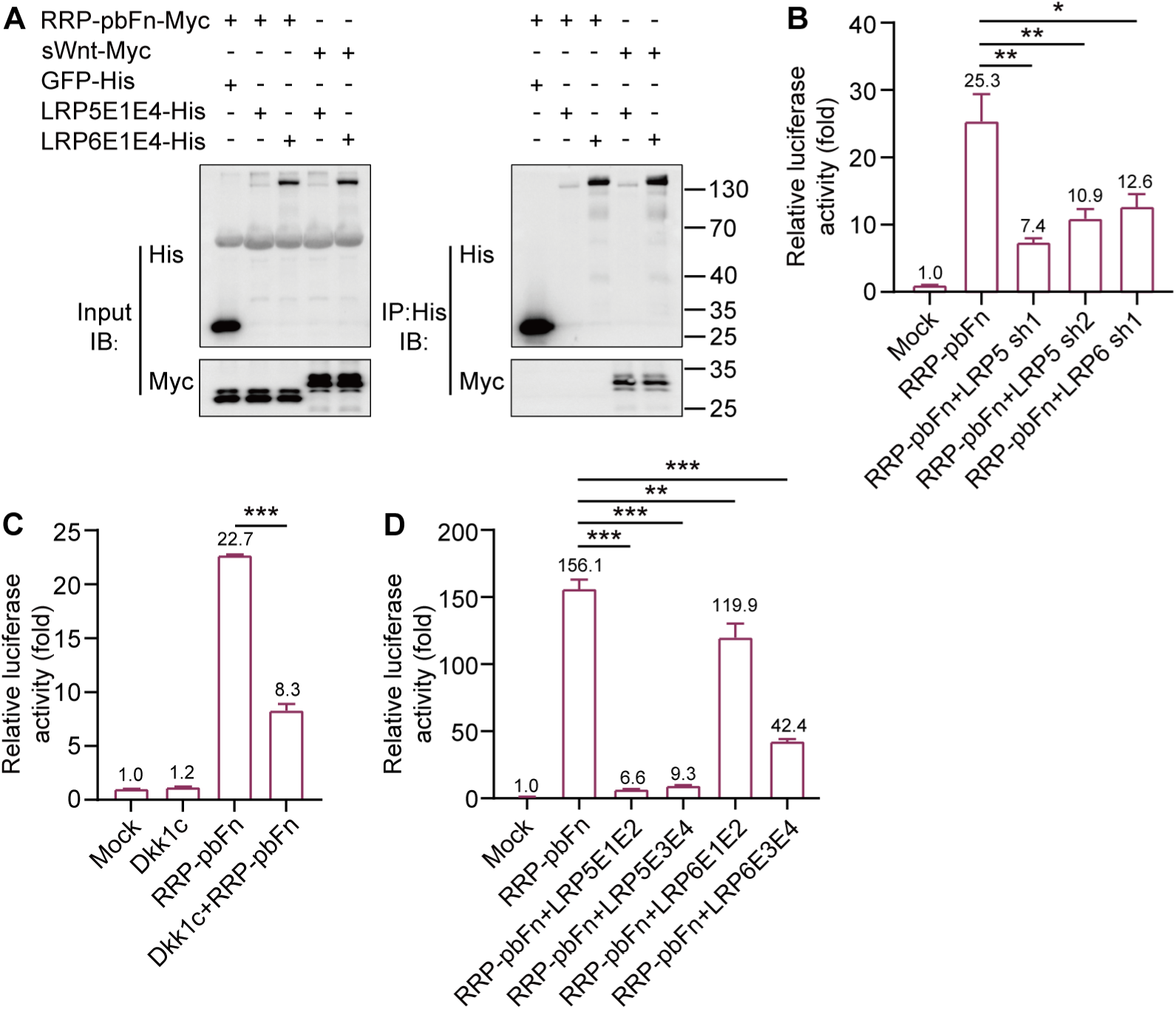
RRP-pbFn does not bind to LRP, but triggers FZD-LRP interaction for Wnt/β-catenin signaling activation. (A) Co-IP analysis of RRP-pbFn and LRP5/6 ectodomain. Supernatant was subjected to immunoprecipitation of His-tagged proteins followed by immunoblotting for anti-Myc. (B) TOP-Flash luciferase reporter assay showing the Wnt activity induced by overexpression of RRP-pbFn with LRP5 sh, LRP6 sh. (C) TOP-Flash luciferase reporter assay showing the Wnt activity induced by overexpression of Dkk1c, RRP-pbFn, or their combination. (D) TOP-Flash luciferase reporter assay showing the impact of L5E1E2, L5E3E4, L6E1E2, or L6E3E4 on RRP-pbFn-mediated activation of Wnt/β-catenin signaling. TOP-Flash data represent mean ± SD; *n* = 3 independent experiments. \**p* < 0.05; \*\**p* < 0.01; \*\*\**p* < 0.001.

To determine whether RRP-pbFn-activated FZD could stimulate Wnt/β-catenin signaling in absence of LRP, short hairpin RNAs (shRNAs) were utilized to suppress the expression of LRP5/6, and knockdown efficiency was confirmed by qRT-PCR analysis (Figures S4A and S4B). It was found that LRP5/6 shRNAs mediated knockdown of LRP5 and LRP6 inhibited RRP-pbFn or sWnt-induced Wnt responses demonstrated by the reduced expression of the TOP-Flash reporter in HEK 293T cells (Figures 4B and S4C). Then, we overexpressed Dkk1c to restrain the function of LRP5/6 in the initiation of Wnt signal transduction, which resulted in abolishing Wnt/β-catenin signaling activation by sWnt and inhibiting RRP-pbFn-induced Wnt/β-catenin signaling activation (Figures 4C and S4D). To further explore the impact of interfering with endogenous transmembrane receptors FZD and LRP5/6 dimerization in Wnt/β-catenin signaling activation, we overexpressed LRP5 ectodomain LRP5E1E2, LRP5E3E4, and LRP6 ectodomain LRP6E1E2, LRP6E3E4. We found that weakening of the interaction between endogenous receptors FZD and LRP5/6 restrained RRP-pbFn or sWnt-induced Wnt/β-catenin signaling activation (Figures 4D and S4E). These findings indicate that the activation of Wnt signaling by RRP-pbFn does not bind to the LRP5/6 receptor, but rather triggers FZD-LRP interaction.

### Generation of high-efficiency LRP antagonist by fusing RRP with Dkk1c

Given that RRP(R52I)-pbFn inhibited the Wnt/β-catenin signaling activation induced by RRP-pbFn (Figure S2B), we proceeded to identify its potential as an antagonist against sWnt-induced Wnt/β-catenin signaling, revealing a modest antagonistic effect (Figure S5). To develop a synthetic antagonist for effectively blocking Wnt ligand or sWnt-dependent Wnt/β-catenin signaling activation, we replaced pbFn with the high-affinity LRP5/6 receptor binder Dkk1 or Dkk1c, considering the partial redundancy of 10 FZD receptors and the binding specificity of pbF to FZD1, 2, 5, 7, and 8.^28, 37^ As expected, both Dkk1 and Dkk1c partially blocked sWnt-induced Wnt responses with or without R-spondin1, while RRP-Dkk1 exhibited further inhibition and RRP-Dkk1c fully suppressed Wnt/β-catenin signaling activation to less than 5% (Figures 5A and 5B), with RRP-Dkk1c demonstrating superior inhibitory activity compared to RRP(R52I)-pbFn (Figure S5). Through mutagenesis of RRP(R52I) and RRP(R55M) (Figure 1K), we unexpectedly observed that RRP(R52I)-Dkk1c and RRP(R55M)-Dkk1c inhibited Wnt/β-catenin signaling activation by sWnt (Figure 5C). These results suggest that RRP played distinct regulatory roles in fusion with pbFn and Dkk1c.

**Figure 5.**
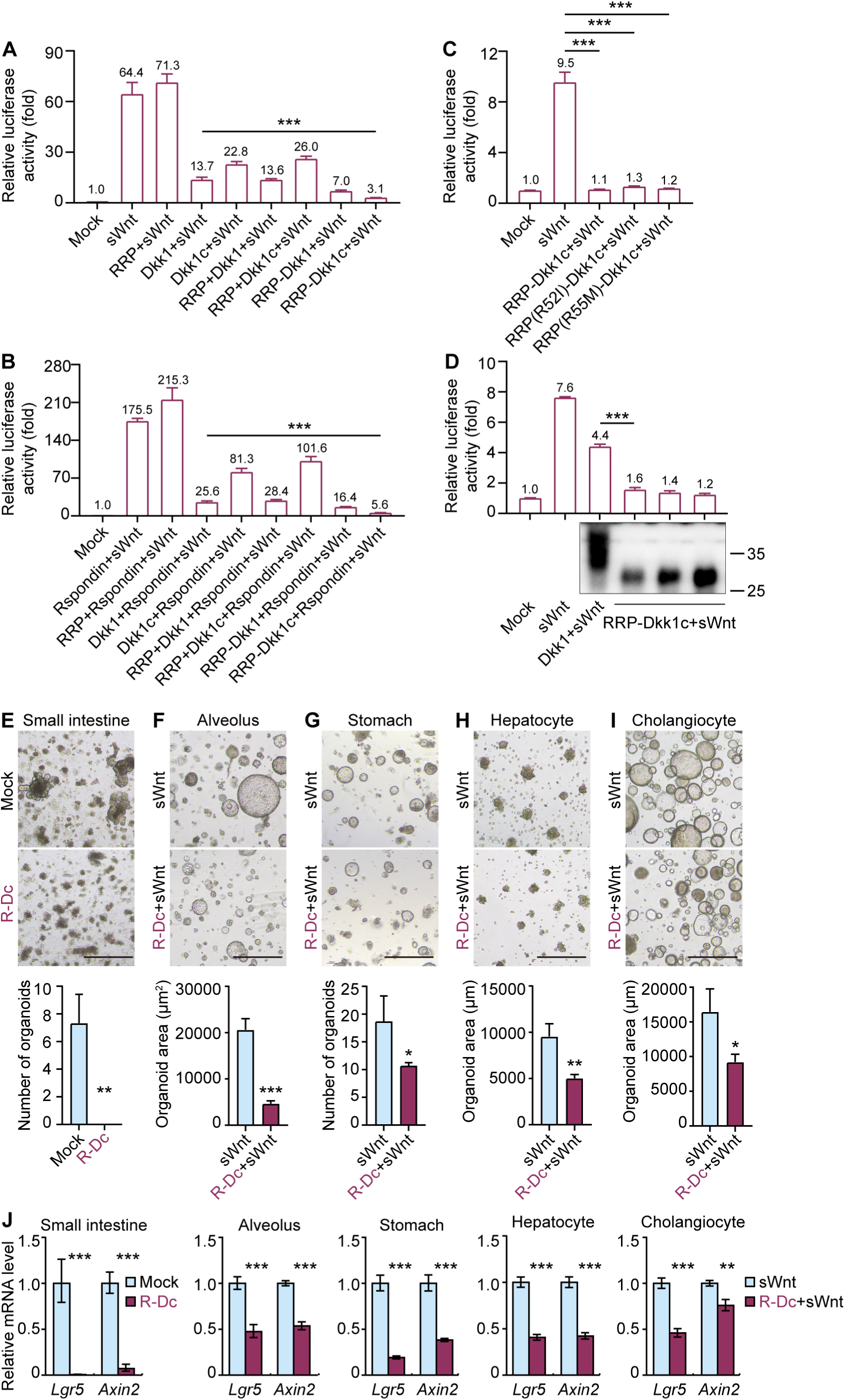
Synthetic LRP antagonist RRP-Dkk1c is a more potent Wnt/β-catenin signaling inhibitor. (A and B) TOP-Flash luciferase reporter assay for evaluating Wnt activity in HEK 293T cells induced by 50 ng/mL sWnt (A) or 50 ng/mL sWnt+250 ng/mL R-spondin1 (B), and in combination with overexpressing the indicated proteins. (C) TOP-Flash luciferase reporter assay for evaluating Wnt activity in HEK 293T cells induced by 50 ng/mL sWnt, and in combination with overexpressing RRP-Dkk1c, RRP(R52I)-Dkk1c, or RRP(R55M)-Dkk1c. (D) TOP-Flash luciferase reporter assay for evaluating Wnt activity induced by 50 ng/mL sWnt, and in combination with overexpressing RRP-Dkk1c or Dkk1. The bottom panel is detection of Dkk1 and RRP-Dkk1c in supernatant after overexpression without sWnt by Western blot for anti-Dkk1. (E) Representative bright-field images of mouse small intestinal organoids expanded in culture media with or without recombinant RRP-Dkk1c. The bottom panel is the quantification of the organoid number. R-Dc: RRP-Dkk1c. (F-I) Representative bright-field images of mouse organoids derived from alveolus (F), stomach (G), and hepatocyte (H) expanded in culture media with 50 ng/mL sWnt, and human cholangiocyte (I) expanded in culture media containing 50 ng/mL sWnt and 3 μM IWP-2, and supplemented with or without recombinant RRP-Dkk1c. The bottom panels are quantification of the organoid number (stomach) or organoid area (alveolus, hepatocyte, and cholangiocyte). The scale bar represents 500 μm. R-Dc: RRP-Dkk1c. (J) qRT-PCR analysis of gene expression of Wnt/β-catenin signaling target genes. R-Dc: RRP-Dkk1c. Data represent mean ± SD, and *n* = 3 for each organoid type. \**p* < 0.05; \*\**p* < 0.01; \*\*\**p* < 0.001.

### RRP-Dkk1c shows a much higher capacity for Wnt/β-catenin signaling inhibition than currently available LRP antagonist

We compared the capacity of Dkk1 with RRP-Dkk1c for Wnt/β-catenin signaling inhibition in a dose-dependent manner using Wnt reporter assays in HEK 293T cells. RRP-Dkk1c showed higher capacity than Dkk1 for inhibiting Wnt/β-catenin signaling activation induced by sWnt (Figure 5D). Additionally, RRP-Dkk1c exhibited a smaller molecular weight compared to Dkk1 (Figure 5D). In conclusion, water-soluble LRP antagonist RRP-Dkk1c exhibited a smaller molecular weight but more potent inhibitory activity than Dkk1.

We then evaluated the function of LRP antagonist RRP-Dkk1c during the growth of organoids. Treatment of mouse intestinal crypt with recombinant RRP-Dkk1c led to a notable reduction in overall organoid viability (Figure 5E). In addition, the application of recombinant RRP-Dkk1c resulted in decreased number of mouse stomach-derived organoids and the size of alveolus-, hepatocyte-, and human cholangiocyte-derived organoids cultured in sWnt-supplemented medium (Figures 5F-5I). Consistently, the mRNA levels of Wnt target genes *Lgr5* and *Axin2* were significantly down-regulated by recombinant RRP-Dkk1c in these organoids (Figure 5J). In conclusion, our findings provide evidence of the potent inhibitory effect of synthetic LRP antagonist RRP-Dkk1c in impeding organoid growths.

### RRP-Dkk1c abolishes the formation of CT26-derived colon cancer xenograft

Kaplan-Meier analysis of 595 colorectal cancer patients from the Human Protein Atlas Dataset revealed that high Dkk1 expression in colon cancer samples (*n*=455) was significantly associated with increased survival rates compared to samples with low Dkk1 expression (*n*=140) (Figure S6), suggesting the inhibitory role of Dkk1 in Wnt/β-catenin signaling is correlated with higher overall survival rate in colorectal cancer patients. To evaluate the potential of RRP-Dkk1c as a tumor growth inhibitor *in vivo*, a xenograft model was established by subcutaneously injecting mice with stable CT26 cells (Figure 6A). CT26, an undifferentiated colon carcinoma cell line lacking the Apc mutation,^38^ was selected as an appropriate model for investigating the effects of RRP-Dkk1c. Lentivirus-infected cells were sorted to generate stable CT26 cells expressing RRP-Dkk1c-Fc or Fc as a control. Notably, the colony formation assay revealed a significant suppression of colony growth in RRP-Dkk1c-Fc stable CT26 cells, with a cell count of 1.88×10^10^ compared to 3.24×10^10^ in control cells after the seventh passage (Figure 6B). In agreement with *in vitro* findings, injection of 5.0×10^5^ control cells into mice resulted in tumor formation, whereas mice injected with RRP-Dkk1c-Fc stable CT26 cells showed no signs of tumor mass (Figures 6C and 6D). These results collectively demonstrate the potent tumor-suppressive properties of RRP-Dkk1c in colon cancer.

**Figure 6.**
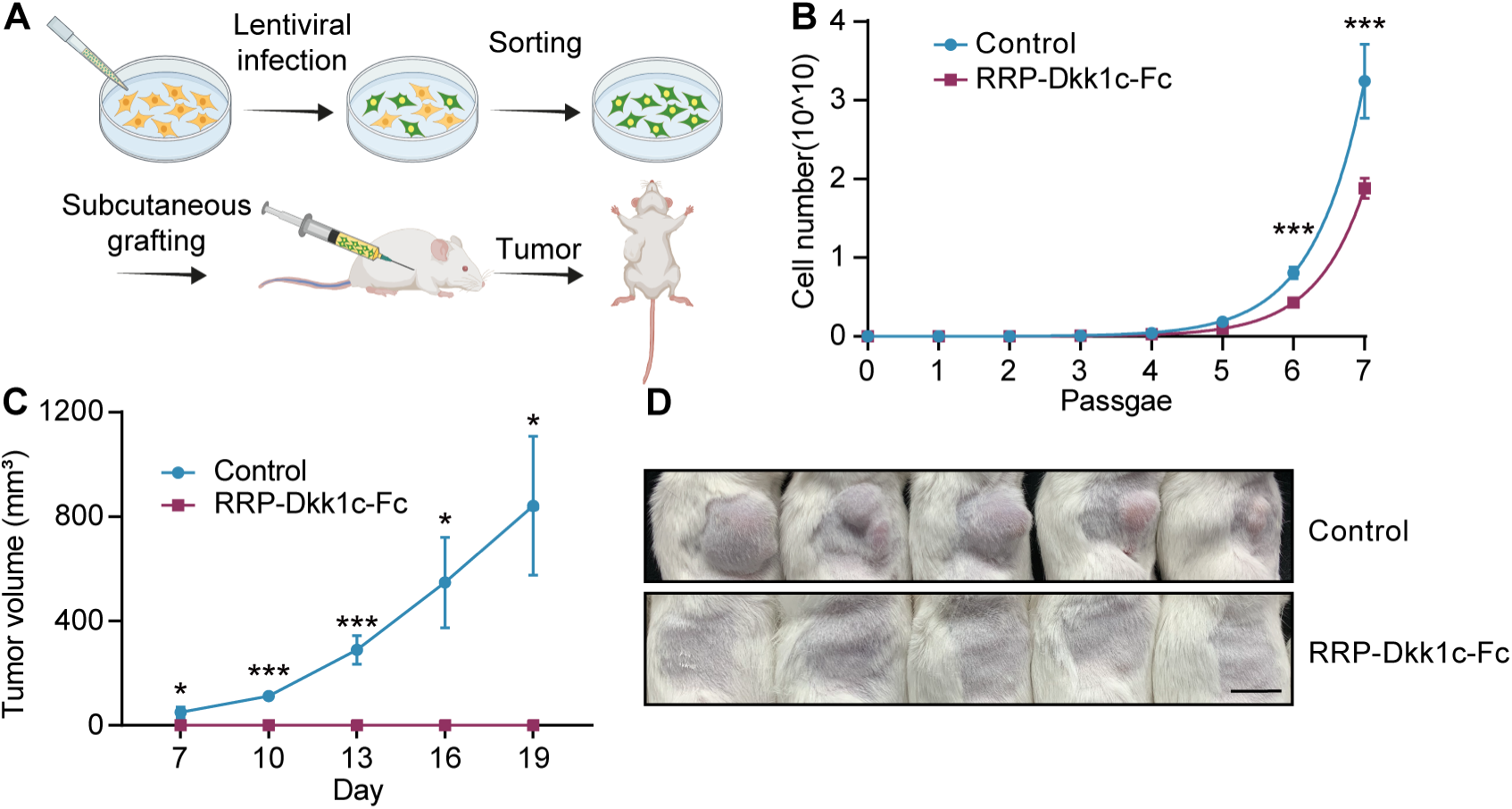
RRP-Dkk1c abolishes the formation of CT26-derived colon cancer xenograft. (A) A scheme depicting the subcutaneous injection of CT26 for the xenograft model. The mice were subcutaneously injected with 5.0×10^5^ sorted CT26 cells. (B) Growth curve of stable CT26 cells expressing RRP-Dkk1c-Fc or Fc (control) from Passage 1 to Passage 7. (C) The volume of subcutaneous xenograft tumors (*n* = 5 mice per group). (D) Photograph of tumors from mice injected with control cells or stable CT26 cells expressing RRP-Dkk1c-Fc on day 19. Data represent mean ± SD, and *n* = 5 mice per group. \**p* < 0.05; \*\**p* < 0.01; \*\*\**p* < 0.001.

## Discussion

The hydrophobicity of Wnt ligands has impeded delineating the molecular mechanisms of Wnt/β-catenin signaling activation and their therapeutic applications. The identification of surrogate Wnt agonist (pbF-Dkk1c) constitutes a remarkable stride in the advancement of research tools.^16^ However, the improved surrogate Wnt (Dkk1c-pbF) exhibits modest efficacy in activating Wnt/β-catenin signaling within our system. Here, we report that the fusion of pMag with pbF recapitulates endogenous ligand activities in the HEK 293T cell line. Specifically, by truncating pMag and pbF, we successfully generate a truncated synthetic FZD agonist known as RRP-pbFn, which demonstrates enhanced potency in comparison to the improved surrogate Wnt. Furthermore, RRP-pbFn, characterized by its lower molecular weight, overcomes the challenges associated with the large molecular size of tetrameric antibody-based Wnt agonists, which comprise two binding sites for FZD and LRP.^17–19^

Due to the remarkable capacity for self-renewal and self-organization, allowing them to closely mimic the structure and function of organs, organoids have emerged as valuable models for studying development and diseases.^7, 9^ We supplement water-soluble recombinant RRP-pbFn to various organoids and observe its effects. Specifically, RRP-pbFn stimulates proliferation in mouse small intestinal organoids, increases the number of mouse stomach organoids, and improves the area of alveolus-, hepatocyte-, and human cholangiocyte-derived organoids. These findings suggest that RRP-pbFn holds the potential to replace Wnt3a-conditioned medium in order to standardize organoid protocols.

In addition, we utilize AAV delivery system to deliver RRP-pbFn *in vivo*. Single treatments with AAV-RRP-pbFn-Fc does not significantly affect liver *GS* expression, but exhibits a significant repression of the periportal marker *Cyp2f2* expression. Notably, when combined with R-spondin1, RRP-pbFn demonstrates a potent regulation of metabolic liver zonation, which is consistent with previous study.^16, 34^ The activity of RRP-pbFn *in vivo* opens up novel therapeutic opportunities for the mobilization of stem cells to facilitate tissue regeneration in human disease contexts characterized by diminished activity.^39, 40^

The typical signaling core at the cell surface comprises a ternary complex of the Wnt ligand, FZD receptor, and LRP receptor.^41^ Notably, our Co-IP data indicates that RRP-pbFn does not bind to the LRP5/6 receptor. We provide evidence that manipulating LRP5/6 expression through LRP5/6 shRNAs, Dkk1c overexpression, and LRP5/6 ectodomain overexpression attenuates RRP-pbFn-induced Wnt/β-catenin signaling activation by weakening the interaction between endogenous FZD and LRP5/6 receptors. These findings suggest that RRP-pbFn does not bind to LRP, but triggers FZD-LRP interaction for Wnt/β-catenin signaling activation. Further investigation is required to elucidate the molecular mechanism underlying the activation of Wnt/β-catenin signaling by RRP-pbFn. Proximity labeling enzymes such as TurboID enable the identification of protein-protein interactions in close proximity by utilizing biotin tagging.^42^ This proteomic approach presents a valuable tool for capturing the interactors of RRP-pbFn at the cell surface, thereby offering novel insights into the molecular mechanisms underlying Wnt/β-catenin signaling activation.

The development and progression of various human cancer types have been associated with enhanced expression of Wnt receptors, attributed to the mutational inactivation of the ubiquitin ligases RNF43/ZNRF3. In this study, we engineer a synthetic water-soluble LRP antagonist by fusing RRP with Dkk1c, which exhibits more potent inhibitory activity compared to Dkk1. Additionally, after subcutaneous injection of mice with stable CT26 cells expressing RRP-Dkk1c-Fc, no discernible tumor masses are observed even 19 days post-injection. These findings highlight the potential of RRP-Dkk1c as a therapeutic strategy for targeting receptor activity in specific subsets of cancer patients.

The varied tertiary structures of agonists and antagonists augment their interactions with specific receptor pockets, granting them with heightened potency and diminished toxicity compared to small molecules.^43^ As a consequence, RRP-pbFn and RRP-Dkk1c exhibit heightened specificity, decreased off-target effects, and enhanced safety profiles.

In summary, we describe two Wnt/β-catenin signaling manipulators. RRP-pbFn serves as a synthetic FZD agonist, overcoming the limitations of natural ligands including high hydrophobicity and low activity. This advancement opens avenues for organoid expansion and the development of disease-related therapeutics. On the other hand, synthetic LRP antagonist RRP-Dkk1c shows promise in inhibiting Wnt/β-catenin signaling for cancer treatment. Collectively, these manipulators hold immense potential for scientific research and a wide range of biomedical applications.

## Methods

### Animals and cell lines

BALB/c mice aged 6-8 weeks were purchased from GemPharmatech. All animal experiments were performed in accordance to protocols approved by the Experimental Animal Ethics Committee of Fudan University. HEK 293T and CT26 cells were cultured in DMEM (Gibco) supplemented with 10 % FBS and 1% Penicillin-Streptomycin at 37 °C in a 5% CO2 environment.

### Plasmids

All constructs, except where indicated, were cloned into pcDNA3.1(+) vector (Thermo Fisher) with an N-terminal tissue plasminogen activator (tPA) signal sequence (MDAMKRGLCCVLLLCGAVFVSA) for overexpression. The sequence of the Vantictumab was retrieved from the published patent, which was subsequently reformatted into pbF. The cDNA of pMag, pbF, and Dkk1c (the C-terminal domain of human Dkk1, residues 177-266) were synthesized by Genscript. The plasmid constructs pMag-pbF and RRP-Dkk1c, designed for secretion of the encoded protein, were generated by cloning pMag or RRP, respectively, along with a flexible linker peptide GSGSG, followed by pbF or Dkk1c, respectively. Plasmids were generated for Co-IP experiment, including constructs encoding RRP-pbFn and sWnt with a C-terminal Myc-tag, as well as constructs encoding LRP5E1E4, LRP6E1E4, and GFP with a C-terminal 6x His-tag. RRP-Dkk1c with a C-terminal Fc-tag was cloned into the lentiviral pLVX-CMV-puro-GFP plasmid. All short hairpin RNA (shRNA) sequences were cloned into the pLKO.1-Puro vector and the sequences are presented in Table S1.

### Conditioned media and protein purification

The pMag-pbF conditioned media was prepared through transient transfection of 293T cells. The medium was replaced with fresh medium 12 hours post-transfection and cells were further incubated for 24 hours. Subsequently, cells were removed by centrifugation. Briefly, for the production of RRP-pbFn and RRP-Dkk1c with a C-terminal Fc-tag recombinant protein, HEK 293T cells were transiently transfected and expression was proceeded for 5 days at 37°C and 8% CO2 with shaking at 125 rpm. After expression, cells were removed by centrifugation and proteins were purified from the conditioned media using rProtein A Sepharose (GE healthcare). The purified proteins were buffer-exchanged into a formulation buffer (20 mM His-HAc, 150 mM NaCl, pH 5.5). Recombinant Fc-tagged RRP-pbFn and Fc-tagged RRP-Dkk1c were abbreviated as recombinant RRP-pbFn and RRP-Dkk1c, respectively.

### TOP-Flash reporter assay

HEK 293T cells were plated into a 48-well cell culture plate and cultured overnight to reach 60-80% confluency. 293T cells were transfected with 50 ng of TOP-Flash plasmid encoding the firefly luciferase and 10 ng of *Renilla* luciferase plasmid per well using VigoFect (Vigorous) according to the manufacturer’s instructions. Twelve hours post-transfection, the medium was changed to fresh medium supplemented with 250 ng/mL R-spondin1, 50 ng/mL sWnt, 50% pMag-pbFn conditioned media, or left untreated for 24 hours as indicated. Cells were lysed with Luciferase Cell Culture Lysis Reagent (Promega) and luciferase activity was measured according to the manufacturer’s protocols. The ratios of firefly luminescence to *Renilla* luminescence were calculated and normalized to the control samples that were left untreated. In terms of illumination, 24 hours after transfection, the cells were illuminated with blue light (470 ± 20 nm, 1.0 mW/cm^2^, 30 seconds pulse every 3 min) for 24 h or kept in darkness.

### Small intestinal organoids

Jejunal tissue (∼10 cm) was isolated, longitudinally cut, and washed with cold DPBS. Subsequently, the villi were carefully scraped off, and tissue was minced and incubated in DPBS containing 5 mM EDTA for 30 min at 4 °C. After transferring the tissue into a tube with fresh DPBS, crypts were released by vigorously shaking then passed through a 70 μm cell strainer (BD). The isolated crypts were pelleted at 600 rpm for 5 min, resuspended in 25 μL Matrigel (Corning), and seeded in a 24-well plate. After polymerization at 37°C for 15 min, culture medium consisting of advanced DMEM/F12 supplemented with N-Acetylcysteine (Sigma-Aldrich), B plus supplement (bioGenous), GlutaMAX (Gibco), 1% Penicillin-Streptomycin (Invitrogen), 50 ng/mL EGF (OrganRegen), 100 ng/mL Noggin (OrganRegen), and 500 ng/mL R-spondin1 (OrganRegen) in the absence or presence of 1 μg/mL RRP-pbFn or 1 μg/mL RRP-Dkk1c was added to each well. Images of the 3D cultured organoids were acquired on Day 5.

### Alveolus organoids

After rinsing the dissected mouse lung with DPBS, incision was made along the edge of the lung lobe. The piece of tissue was subjected to a freshly prepared digestion solution consisting of advanced DMEM/F12 supplemented with 10 μM Y-27632 (Sigma-Aldrich), 400 U/mL collagenase NB 4 (Nordmark), and 10 U/mL Dnase I (Roche) for 50 min at 37°C. The digestion solution was pipetted up and down and then filtered through a 70 μm cell strainer. After centrifugation of the filtrate at 350 g for 3 min, the supernatant was discarded, and Red Blood Cell Lysis Buffer (eBioscience) was added for 3 min followed by DPBS washing. The mixture was centrifuged and then embedded in Matrigel. The culture medium consisted of advanced DMEM/F12 supplemented with B plus, GlutaMAX, N-Acetylcysteine, 1% Penicillin-Streptomycin, 10 mM HEPES (Gibco), 5 mM Nicotinamide (Sigma-Aldrich), 0.5 μM A83-01 (Tocris), 10 μM Y-27632, 0.5 μM SB-202190 (Sigma-Aldrich), 50 ng/mL EGF, 500 ng/mL R-spondin1, 100 ng/mL Noggin, 100 ng/mL FGF10 (OrganRegen), and 25 ng/mL FGF7 (OrganRegen) in the absence or presence of 50 ng/mL sWnt (OrganRegen), 1 μg/mL RRP-pbFn or 1 μg/mL RRP-Dkk1c. Images of the 3D cultured organoids were acquired on Day 5.

### Gastric organoids

Gastric glands units were isolated from mouse pyloric stomach as previously described with some modifications.^44^ Briefly, the stomach was opened along the greater curvature, and the pylorus was isolated followed by DPBS washing. The muscular layer of the stomach was removed and the remaining epithelia was divided into 5 mm pieces and incubated in digestion solution consisting of advanced DMEM/F12 supplemented with 2 mg/mL collagenase type I (Thermo), 100 μg/mL primocin (Invivogen), 10% FBS, and 10 U/mL DNase I for 20-30 min at 37°C. The isolated gastric glands were filtered through a 70 μm cell strainer then a total of 200 glands mixed with Matrigel were plated in 24-well plate. Gastric culture medium consisted of advanced DMEM/F12 supplemented with B plus, GlutaMAX, N-Acetylcysteine, 1% Penicillin-Streptomycin, Y-27632, 50 ng/mL EGF, 500 ng/mL R-spondin1, 100 ng/mL Noggin, 100 ng/mL FGF10, 10 nM Gastrin I (Sigma-Aldrich), and 100 μg/mL Primocin in the absence or presence of 50 ng/mL sWnt, 1 μg/mL RRP-pbFn or 1 μg/mL RRP-Dkk1c. Images of the 3D cultured organoids were acquired on Day 4.

### Hepatocyte organoids

Primary hepatocytes were isolated from mice as previously described.^45^ Briefly, after placing catheter into the portal vein, the inferior vena cava was cut and the liver was perfused with pre-warmed perfusion medium. Then, perfusion was performed with pre-warmed digestion medium including collagenase type IV and Ca^2+^ for 3-5 min. After dissociation, cells were filtered through a 70 μm cell strainer. Hepatocytes were further separated and purified by centrifugation at 50 g for 3 min and percoll gradient centrifugation was performed. In a 24-well plate, 5×10^4^ isolated hepatocytes mixed with Matrigel at a concentration of 75% diluted by culture medium were used per well. Culture medium was composed of advanced DMEM/F12 supplemented with B plus, GlutaMAX, N-Acetylcysteine, 1% Penicillin-Streptomycin, HEPES, Nicotinamide, 1 μM A83-01, Y-27632, 50 ng/mL EGF, 500 ng/mL R-spondin1, 100 ng/mL FGF10, 10 nM Gastrin I, 25 ng/ mL HGF (OrganRegen), 100 ng/mL TGF-α (R&D Systems), and 100 ng/mL TNF-α (R&D Systems) in the absence or presence of 50 ng/mL sWnt, 1 μg/mL RRP-pbFn or 1 μg/mL RRP-Dkk1c. Images of the 3D cultured organoids were acquired on Day 4.

### Human cholangiocyte organoids

Cholangiocyte organoids from liver biopsies obtained from donor livers were generated as previously described.^46^ Briefly, Biopsies were minced and digested by incubation in advanced DMEM/F12 supplemented with 2 mg/mL collagenase type I and 10 U/mL DNase I for 20-30 min at 37°C. Digestion solution was diluted by adding cold advanced DMEM/F12 supplemented with 1% Penicillin/Streptomycin and 1% FBS. The cell suspension was filtered through a 70 μm cell strainer and centrifuged at 400 g, 4°C for 3 min. Supernatant was removed, and the cell pellet was suspended with Matrigel which was allowed to solidify for 15 min at 37°C before cholangiocyte organoid culture medium was added. Culture medium was composed of advanced DMEM/F12 supplemented with B plus, GlutaMAX, N-Acetylcysteine, Penicillin-Streptomycin, HEPES, Nicotinamide, 2 μM A83-01, 50 ng/mL EGF, 500 ng/mL R-spondin1, 100 ng/mL FGF10, 10 nM Gastrin I, 25 ng/mL HGF, and 10 μM Forskolin (Sigma-Aldrich). After passaging cholangiocyte organoids, 3 μM IWP-2 was added in culture medium to inhibit endogenous Wnt lipidation and secretion in the absence or presence of 50 ng/mL sWnt, 1 μg/mL RRP-pbFn or 1 μg/mL RRP-Dkk1c. Images of the 3D cultured organoids were acquired on Day 5.

### Immunoprecipitation and immunoblotting

Secreted RRP-pbFn-Myc, sWnt-Myc, GFP-His, LRP5E1E4-His, and LRP6E1E4-His were overexpressed in HEK 293T cells. After expression, cells were removed by centrifugation and supernatant was mixed with 1:1 as indicated, then HEK 293T cells were overlaid with the mixture for 60 min at 37°C. The mixture was used for immunoprecipitation with Ni Sepharose excel (GE healthcare) according to the manufacturer’s protocols. Immunoblotting was performed using the following antibodies: α-His (Proteintech, 1:3000), α-Myc (Abclonal, 1:5000).

### Immunofluorescence

The small intestine and liver of mouse were fixed overnight in 4% paraformaldehyde at 4°C and processed for paraffin embedding. Subsequently, 5-μm sections were stained with the following antibodies following citrate antigen retrieval and blocking with 10% normal donkey serum: mouse anti-glutamine synthetase (BD Bioscience, 1:2000), mouse anti-Ki67 (BD, 1:500). Then sections were incubated with Alexa Fluor 488 (Jackson ImmunoResearch, 1:200) and Cy3 (Jackson ImmunoResearch, 1:200)-labeled secondary antibodies, counterstained with DAPI at room temperature in the dark. Images were captured with Olympus FV3000 Confocal Laser Scanning Microscope.

### RNA extraction and quantitative Real-Time PCR

Total RNA was extracted using RNAprep Pure Micro Kit (Tiangen Biotech) according to manufacturer’s protocols. Purified RNA was reverse-transcribed into cDNAs with GoScript Reverse Transcription System (Promega) according to the manufacturer’s instruction. Quantitative Real-Time PCR was performed using SYBR Green PCR Master Mix (Bimake) and detected by CFX384 Touch System (Bio-rad). Expression levels were normalized to the reference gene *Gapdh*, and the primers were listed in Table S2.

### *In vivo* experiments

A single intravenous injection of 3.0×10^11^ vg/mouse AAV-8 expressing Fc (control) or RRP-pbFn-Fc was administered. After 10 days injection of AAV-8, mice were injected intravenously with 200 µg/mouse recombinant R-spondin1 or PBS daily for 6 days. Intestinal tissues and livers were collected 24 h after the last R-spondin1 injection (*n* = 3 mice per group).

### Mouse xenograft model

CT26 tumor cells infected with either RRP-Dkk1c-Fc or Fc (control) lentivirus were sorted for GFP expression, 5.0×10^5^ sorted GFP^+^ cells were suspended in 125 μL phosphate buffered saline and subcutaneously injected into the right flanks of BALB/c mice. Tumor growth was measured with vernier calipers and tumor volumes were calculated using the modified ellipsoid formula (0.5 × length × width × width).

### Colony formation assay

1.0×10^6^ sorted GFP^+^ CT26 tumor cells were seeded per 6-well plate and cultured in medium until they reached 80–90% confluency. CT26 were then dissociated into single cells using 0.25% Trypsin and the number of cells was counted. The counted cells were considered as passage 1, and 1.0×10^6^ cells were reseeded per 6-well plate for subsequent passages. This process was repeated for a total of seven passages, and each experiment was performed with 5 replicates.

### Survival analysis

The clinical data was obtained from Human Protein Atlas Dataset, and a total of 595 colorectal cancer patients were included for survival analysis. Dkk1 expression levels were determined using FPKM (fragments per kilobase of transcript per million fragments mapped) values, with FPKM > 0 considered as high Dkk1 expression and FPKM = 0 considered as low Dkk1 expression.

### Protein structure modeling

The protein sequences in FASTA format were supplied for SWISS-MODEL.^47^ Subsequently, 3hjk.1 was utilized to identify the optimal template for protein modeling. Through aligning the target sequence with the template, conserved sequences were identified. Based on the homology information, full-atom protein models were generated. The visualization settings of the model were adjusted using ChimeraX.

### Quantification and statistical analysis

Statistical analyses were performed using GraphPad Prism 8. Student’s t-test or one-way ANOVA test was employed to analyze the parametric experimental results, and Log-rank test was used for survival analysis. The mean and standard deviation (SD) or standard error (SE) were reported in the figure legends. A p-value < 0.05 was considered statistically significant.

## Acknowledgments

This work was supported by grants from the National Natural Science Foundation of China (32022022 and 31970761), the Shanghai Pilot Program for Basic Research (Fudan University 2100100-213), the Key Research and Development Program of Yunnan Province (202302AA310024), and the Faculty Resources Project of College of Life Sciences (Inner Mongolia University 2022-103).

## Author Contributions

Q.D., C.N., and B.Z. designed the experiments; Q.D., J.W., Z.L., D.Y., H.Y., J.Z., X.L., J.Z., and H.H. performed the experiments; Q.D., C.N., and B.Z. analyzed the data; B.Z. supervised the work; Q.D. and B.Z. wrote the paper.

## Supplementary Information

**Figure S1.**
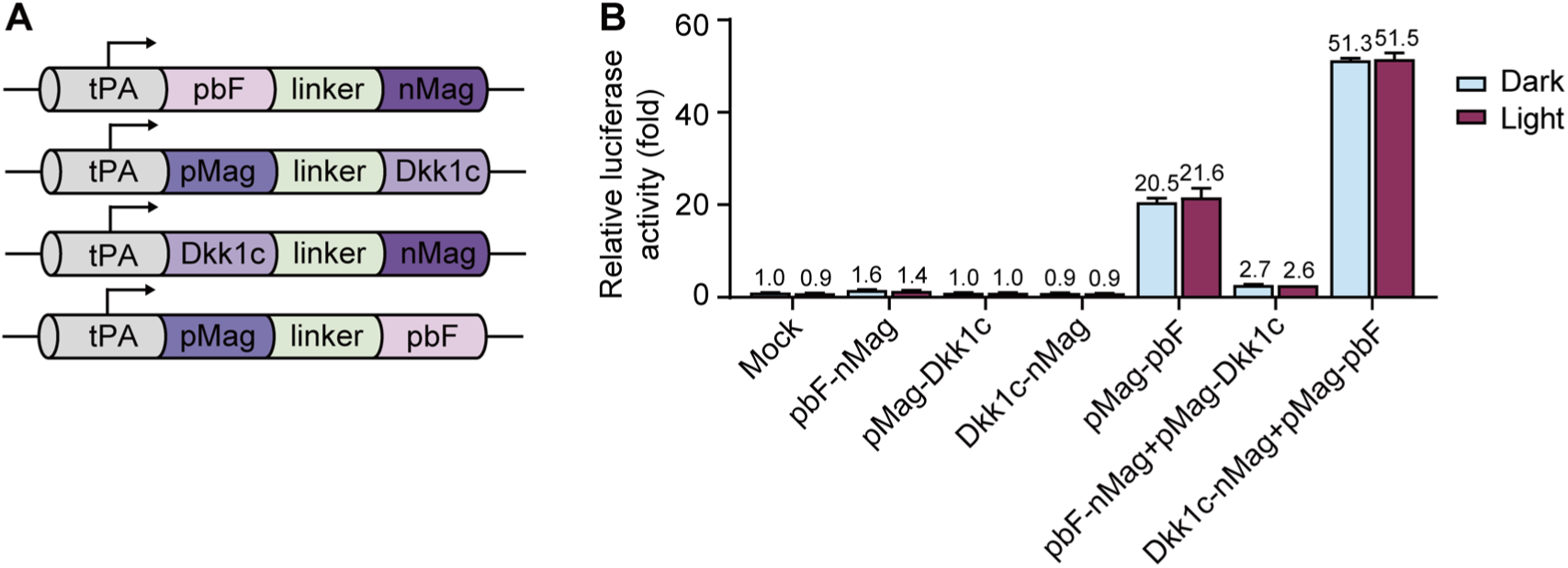
pMag-pbF activates Wnt/β-catenin signaling without blue light illumination. (A) Scheme depicting combinations of variant sWnt fragments (pbF/Dkk1c) with the pMag/nMag. (B) Luciferase reporter assay for evaluating Wnt activity in HEK 293T cells overexpressing pbF-nMag, pMag-Dkk1c, Dkk1c-nMag, pMag-pbF, and their combinations, with or without illumination. Data represent mean ± SD; *n* = 3 independent experiments.

**Figure S2.**
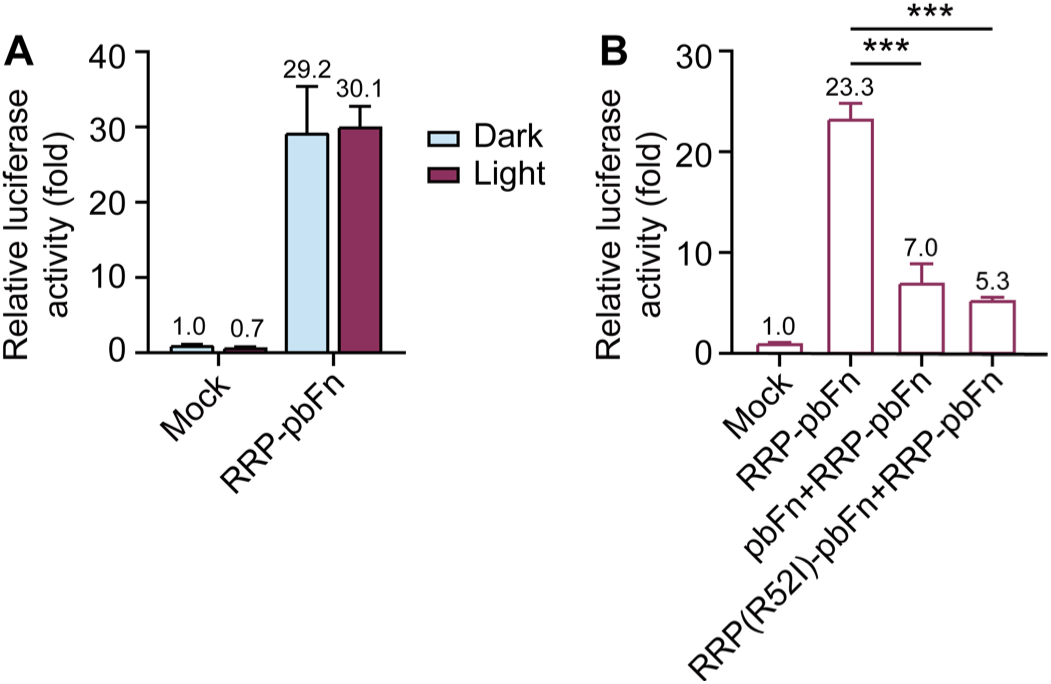
RRP(R52I)-pbFn blocks the Wnt signaling activation induced by RRP-pbFn. (A) Luciferase reporter assay for evaluating Wnt activity in HEK 293T cells overexpressing RRP-pbFn with or without illumination. (B) Luciferase reporter assay for evaluating Wnt activity in HEK 293T cells overexpressing RRP-pbFn, and combinations of RRP-pbFn with pbFn or RRP(R52I)-pbFn, respectively. TOP-Flash data represent mean ± SD; *n* = 3 independent experiments. \**p* < 0.05; \*\**p* < 0.01; \*\*\**p* < 0.001.

**Figure S3.**
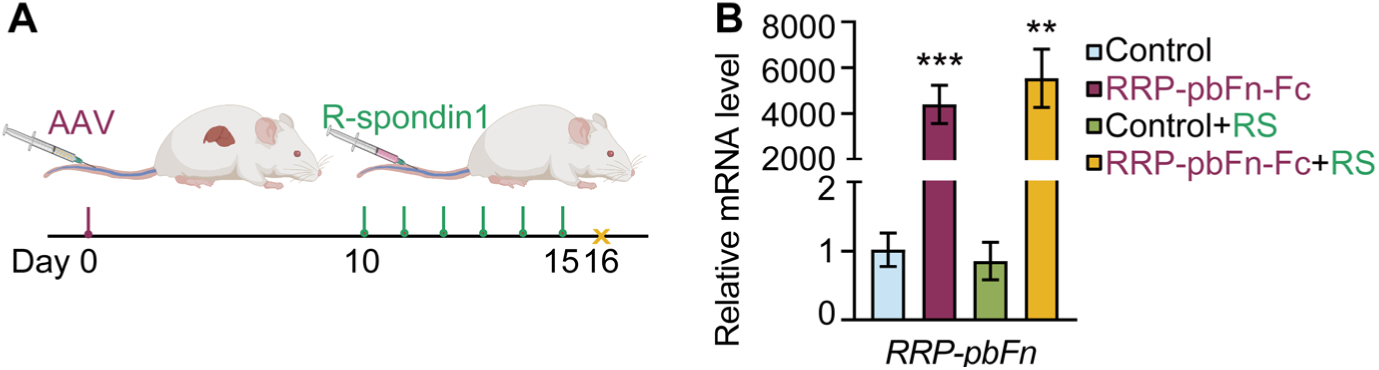
AAV-directed expression of RRP-pbFn-Fc in mice. (A) Scheme depicting intravenous injection of AAV and R-spondin1. (B) qRT-PCR validation of AAV-mediated transgene RRP-pbFn-Fc expression in mouse liver. RS: R-spondin1. Data represent mean ± SD. *n* = 3 mice per group. \**p* < 0.05; \*\**p* < 0.01; \*\*\**p* < 0.001.

**Figure S4.**
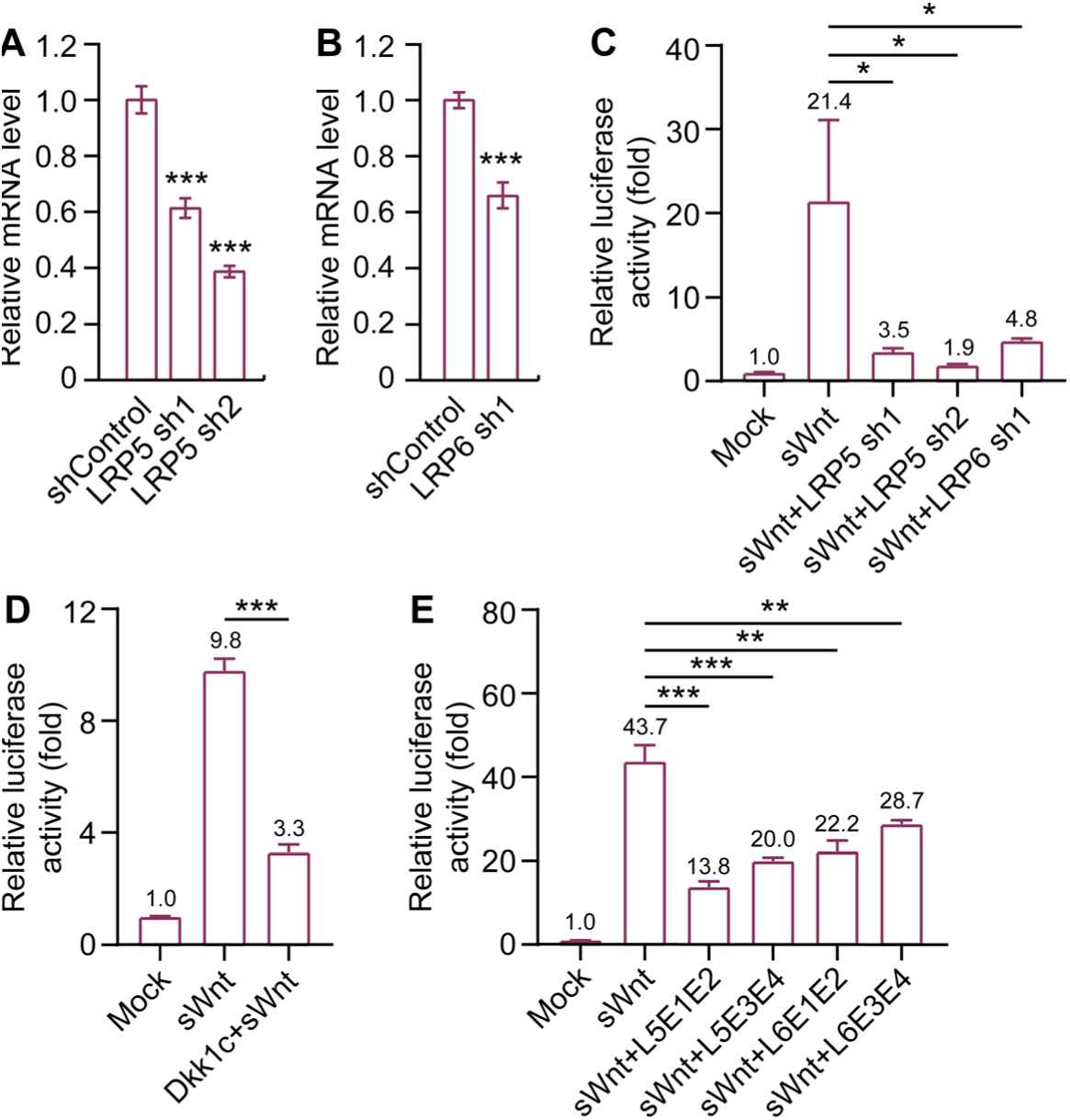
Blocking the function of LRP5/6 prevents sWnt-induced Wnt signaling. (A and B) qRT-PCR analysis of the expression of *LRP5* (A) or *LRP6* (B) in HEK 293T cells after interference with LRP5 shRNAs, LRP6 shRNA, or scrambled shRNA (shControl). (C) TOP-Flash luciferase reporter assay showing the Wnt activity induced by 50 ng/mL sWnt alone and in combination with overexpressing the indicated shRNAs. (D) TOP-Flash luciferase reporter assay showing the Wnt activity induced by 50 ng/mL sWnt alone and in combination with overexpression of Dkk1c. (E) TOP-Flash luciferase reporter assay showing the impact of L5E1E2, L5E3E4, L6E1E2, or L6E3E4 on 50 ng/mL sWnt mediated activation of Wnt signaling. TOP-Flash data represent mean ± SD; *n* = 3 independent experiments. \**p* < 0.05; \*\**p* < 0.01; \*\*\**p* < 0.001.

**Figure S5.**
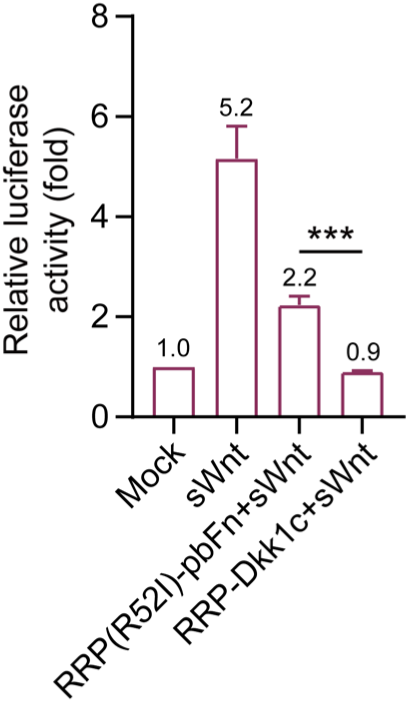
RRP-Dkk1c blocks the Wnt signaling activation induced by sWnt. TOP-Flash luciferase reporter assay for evaluating Wnt activity in HEK 293T cells induced by 50 ng/mL sWnt alone and in combination with overexpressing RRP(R52I)-pbFn or RRP-Dkk1c. Data represent mean ± SD; *n* = 3 independent experiments. \**p* < 0.05; \*\**p* < 0.01; \*\*\**p* < 0.001.

**Figure S6.**
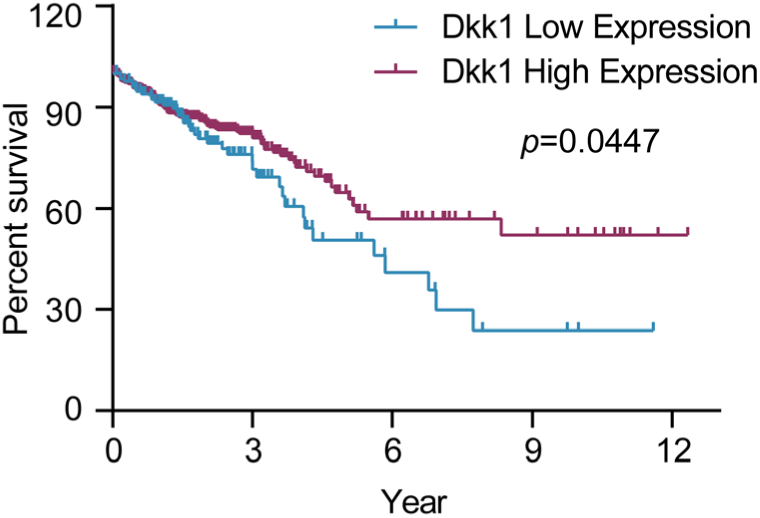
Kaplan-Meier estimates survival probability by Dkk1 expression level in colorectal cancer. Overall survival rate of colorectal cancer patients according to different expression level of Dkk1. The endpoint was 13 years. Survival curves were compared via Log-rank test and visualized by Kaplan-Meier method.

**Table S1.**
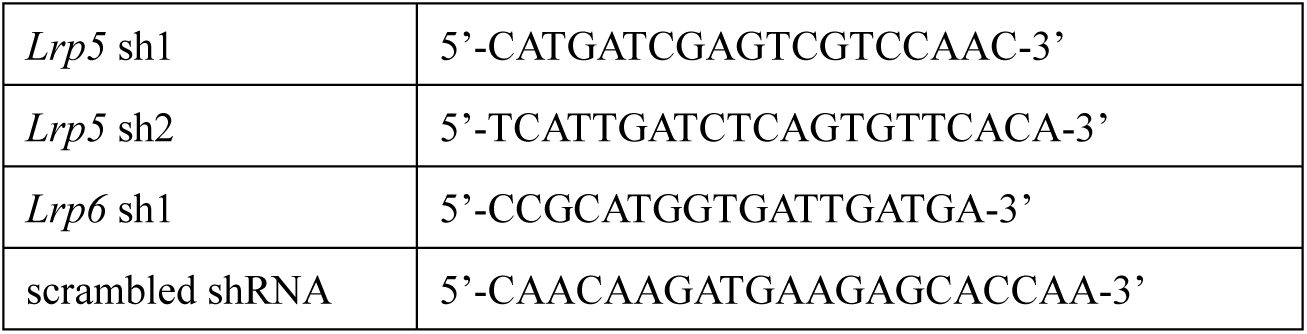
Primers for shRNA.

**Table S2.**
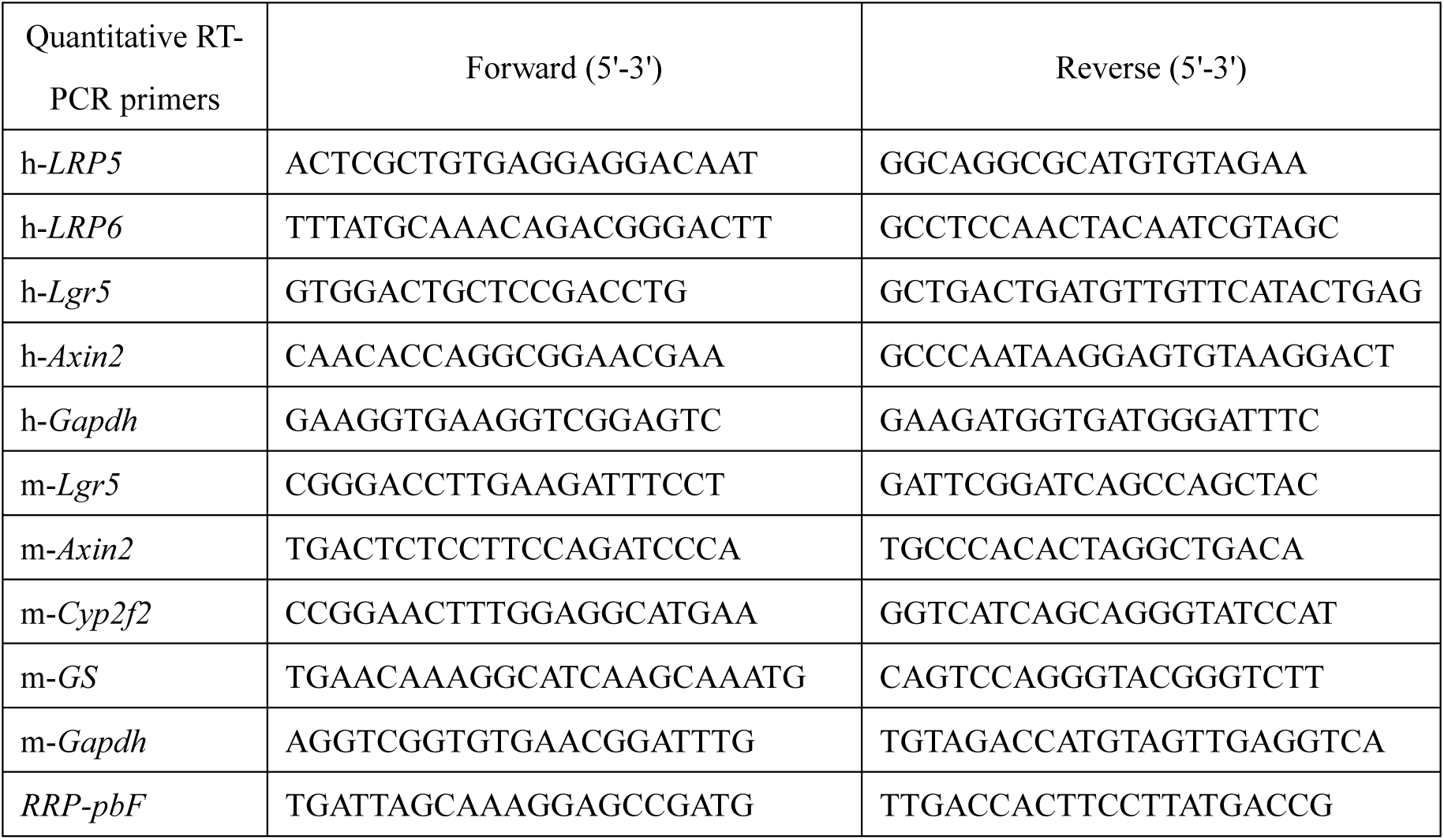
Primers for RT-qPCR.

